# Genome-wide assays to characterize rAAV integration into human genomic DNA in vivo

**DOI:** 10.1101/2023.08.22.554338

**Authors:** Jaime Prout, Jessica Von Stetina, Gustavo Cerqueira, Madison Chasse, Huei-Mei Chen, Andrew Pla, Rachel Resendes, Danielle Sookiasian, Cagdas Tazearslan, Jason Wright, John F. Thompson

## Abstract

Adeno-associated viral (AAV) vectors are used to treat genetic diseases, expressing therapeutic genes from both extrachromosomal episomes and payloads that integrate into the host genome. Assays were developed to evaluate HR-mediated on-target integration and the potential occurrence of off-target integration. While many studies have addressed elements of these processes, proper characterization requires long-read sequencing to ensure that integrated viral DNA is examined and not the more prevalent episomes. We used Oxford Nanopore to characterize integrated DNA and scan the whole genome for off-target integrations. These assays were applied to cell-based and in vivo models to study vectors that correct phenylketonuria (PKU), caused by loss of phenylalanine hydroxylase (*PAH*). Administration of the human-specific vector in a humanized-liver mouse xenograft model resulted in stable, nuclease-free integration into *PAH*. Because detection of rare integration events in a much larger pool of episomal DNA is subject to artifacts, careful assay validation was required. A long-read, genome-wide assay capable of detecting on- and off-target vector integrations showed no evidence of off-target integration. Artifactual false positive events were below the limit of blank. These data support rAAV as an investigational therapeutic for genetic diseases and reinforce the need for characterization of integration assays to avoid artifacts.

## Introduction

rAAV DNA has been shown to exist in cells both chromosomally after genome integration and extra-chromosomally as an episome.^1–4^ Upon entry into the cell’s nucleus, the single-stranded rAAV vector genomes can either integrate into the genome via homologous recombination (HR) or be processed into extra-chromosomal episomes. If recombining, the two homology arms containing the homologous sequences integrate into the target genomic DNA, resulting in the insertion of the vector payload DNA located between the two homology arms. Non-homology-driven integration of AAV has been extensively studied at locations such as the AAVS1 site on chromosome 19.^3,5^ In contrast, HR-dependent rAAV integration can occur when there is homologous sequence shared between the rAAV vector and the genome, resulting in HR-driven, site-specific integration. For example, recent work using rAAV-mediated HR integration into the albumin locus took advantage of both the fidelity of HR integration and the high expression of albumin to drive transcription of the desired gene.^6–8^

Evidence for the integration of AAV and other viruses into the human genome has been demonstrated using a succession of increasingly robust methodologies. Wild-type AAV was found to integrate specifically into chr19 using in situ hybridization.^3,4^ This finding was followed by the detection of AAV and other viral integrations by FISH,^9^ plasmid rescue,^10^ LAM-PCR,^11,12^ nrLAM-PCR,^13^ brute force NGS,^14^ targeted capture,^15,16^ single-cell approaches,^17,18^ and a comparison of NGS-based approaches.^19^ Some methods can be used across a broad range of viruses while others have significant limitations in particular situations. Methods that work well with efficiently recombining systems like retroviruses may not be feasible with less efficient systems like AAV where the high background of non-integrated viral genomes can lead to many artifacts. In addition, information gathered from each assay varies. No sequence information is generated with qPCR,^20^ while other methods can produce integration junction sequence^21^ or full-length integrated viral sequence data.

Cells transduced with rAAV can have hundreds to thousands of copies of episomal DNA per cell. The episomal and integrated DNA contain long stretches that are identical to each other in both the homology arms and payload. Because of the significant mismatch in copy number, episomal DNA can contaminate reads purportedly from integrated DNA. To ensure that homology arms from integrated viruses are evaluated rather than the more common episomal DNA, each sequence read should span the viral payload through the homology arm and into the unique genomic sequence adjacent to the integration event. Previous reports using short-read sequencing technology showed no additional changes to the genome,^22^ but this finding requires confirmation via long-read sequencing. Additionally, multiple classes of other artifacts potentially arise during sample processing and NGS library preparation and these must be addressed when determining assay sensitivity.^23^ Appropriate controls need to be included for an assay to be meaningful.

Accurate assessment of on- and off-target integration is essential because rAAV integration has been proposed as a means of achieving stable correction of genetic defects. Episomal expression of a vector containing a codon-optimized human PAH (CO-hPAH) DNA driven by a liver-specific promoter results in episomal expression and correction of phenylketonuria (PKU) in treated mice.^24^ Subsequent studies showed that CO-hPAH with homology arms packaged in AAVHSC15 could induce nuclease-free, HR-mediated integration into the human *PAH* locus. These ∼900 bp homology arms are locus- and species-specific and were designed to minimize potential off-target HR integration. To better characterize HR-driven rAAV integration, we have developed long-read sequencing assays to characterize all integration sites, independent of genomic location. These methods use techniques that have not been previously applied to rAAV integration.

## Results

### rAAV vectors and on-target integration

Assays to assess on-target integration via ddPCR and sequencing short PCR amplicons have been described previously.^22^ ddPCR is useful when the number of episomes is low, but it becomes impractical when high doses of rAAV are administered and the ratio of episomal to integrated DNA becomes excessive. The large number of episome-only droplets leads to an episomal copy number that overwhelms the integration signal. When ddPCR could not be used, amplicon sequencing was carried out using short-read sequencing across the integration junctions. Because the sequences of both the integrated vector DNA and adjacent genomic DNA are known, it is straightforward to design PCR primers that allow selective amplification of the wild-type genomic DNA or the integrated DNA. However, the large excess of episomal DNA and the potential presence of concatemers of variable structure^25^ can introduce artifacts during sample preparation and amplification. To minimize this, the amplicons must be sufficiently long that there is payload DNA sequenced on one end of the homology arm and a non-viral genomic DNA sequence on the other end. When quantitation is desired, amplification conditions also need to be chosen such that a high level of episomal DNAs present does not differentially affect the integrated/non-integrated DNA ratio. This is most easily done by making the primer required to amplify both these DNAs product-limiting. With these requirements, primers have been created that allow selective amplification and quantitation of both wild-type and integrated DNAs.

Because rAAV integration is homology-dependent, different vectors must be used for integration into mouse or human genomic DNA. Integration is more efficient in vivo than in vitro so special models must be used to examine integration into human DNA. A relevant system for assessing rAAV integration behavior is human hepatocytes that are engrafted into FRG® mouse livers.^26^ These mice have defective immune systems that allow the engraftment and growth of human cells. The mouse hepatocytes are engineered to be sensitive to a chemical challenge. In the appropriate conditions, most mouse hepatocytes die and are replaced by human donor hepatocytes. While this model provides an in vivo setting, the mice are not robust due to their defective immune system and do not survive as long as normal mice. This makes them much more difficult to maintain. The challenges in working with this model limit the experiments that can be done; but, despite the limitations, the model provides insight into how the vector is likely to behave when administered to humans.

Three different vectors have been used for the integration experiments described here, two with human homology arms but different payloads, and one with no homology arms. Vectors are named by the species/presence of homology arms (hHA), whether a promoter is present (LP1), and the protein product (hPAH). Some of these vectors have been described previously^22^ and are illustrated in Supplemental Figure 1. Mice are injected with either Formulation Buffer (FB) as a control or differing concentrations of species-specific vectors at varying concentrations. Figure 1 shows 3-7% integration can be achieved in human hepatocytes with human homology arms as measured using the long-read PCR amplicon method previously described.^22^ A common genomic primer is amplified with either a payload or genomic primer located beyond the homology arms and the ∼1.4 kb amplicons are long-read sequenced using Oxford Nanopore Technologies (ONT). The read counts of integrated DNA versus wildtype DNA are then compared. The amount of integration is dose dependent. The high level of episomal vector genomes (VGs) in the vector-treated samples prevents the ddPCR method from being used.

**Figure 1:**
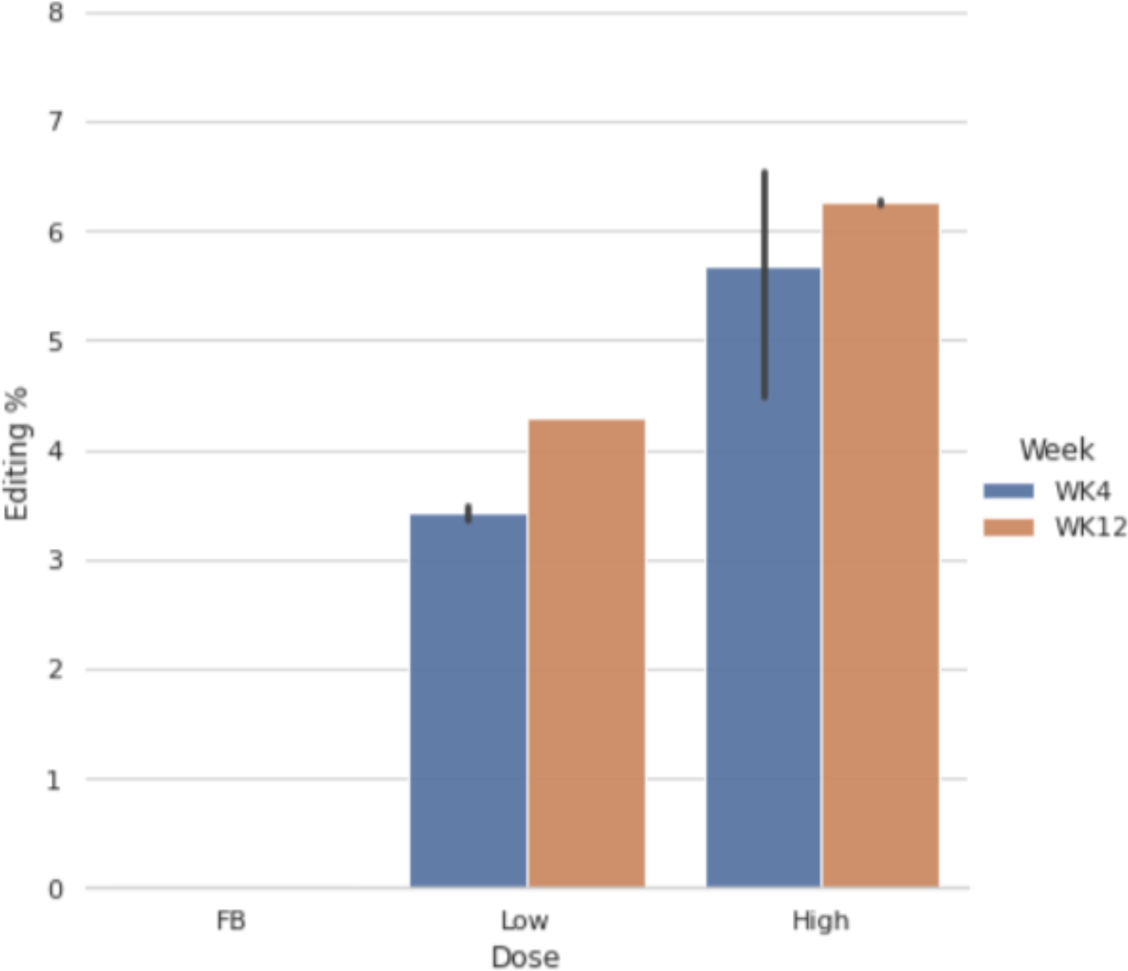
Integration of hPAH into human PAH locus. FRG mice were treated with formulation buffer (FB), 7E+13 VG/kg (Low), or 2E+14 VG/kg (High) of hHA-LP1-hPAH. After either 4 (blue) or 12 weeks (brown), mice were sacrificed, and livers removed for isolation of human hepatocytes and DNA analysis.

### Predicted Off-target integrations

If rAAV vectors were to integrate at genomic sites other than their intended targets, it is most likely that such off-target integrations would be directed toward homologous sequences. During vector design, homology arms were chosen to have minimal sequence identity with the rest of the genome. As a result, all sequences elsewhere in the genome are poor matches for the hHA-LP1-hPAH homology arms. To test the most likely of the poor matches, predicted off-target integration sites were selected based on two criteria, a minimum sequence length of 35 bp and a minimum sequence identity of 60% relative to any sequence in the homology arms. Only six genomic regions could be identified that met both these criteria. The six identified regions had 38-52 bp homology with 60-85% identity relative to the hHA-LP1-hPAH homology arms. Because the sequence of both the homology arms and the predicted integration site is known, it is straightforward to design primers that allow specific amplification for putative off-target integration sites at 5 of these 6 regions (Supplemental Table 1).

To generate positive control DNAs for the potential integration sites, gBlock DNA was synthesized that contained a sequence corresponding to what the putative integration sites would have if the junctions were present in the genome. These were used to spike into control human genomic DNA at molar ratios starting at 1:10 and serially diluting 10x down to 1:10,000. Each spiked, positive-control integration site could be detected at all ratios when amplified with the integration-specific primers. This established the limit of detection as 1 in 10,000. When genomic DNA from the FRG mice treated with hHA-LP1-hPAH was tested, no signal at any of the predicted off-target sites could be detected (Figure 2), indicating that the five tested sites were not subject to off-target integration at rates above the Limit of Detection (LoD).

**Figure 2:**
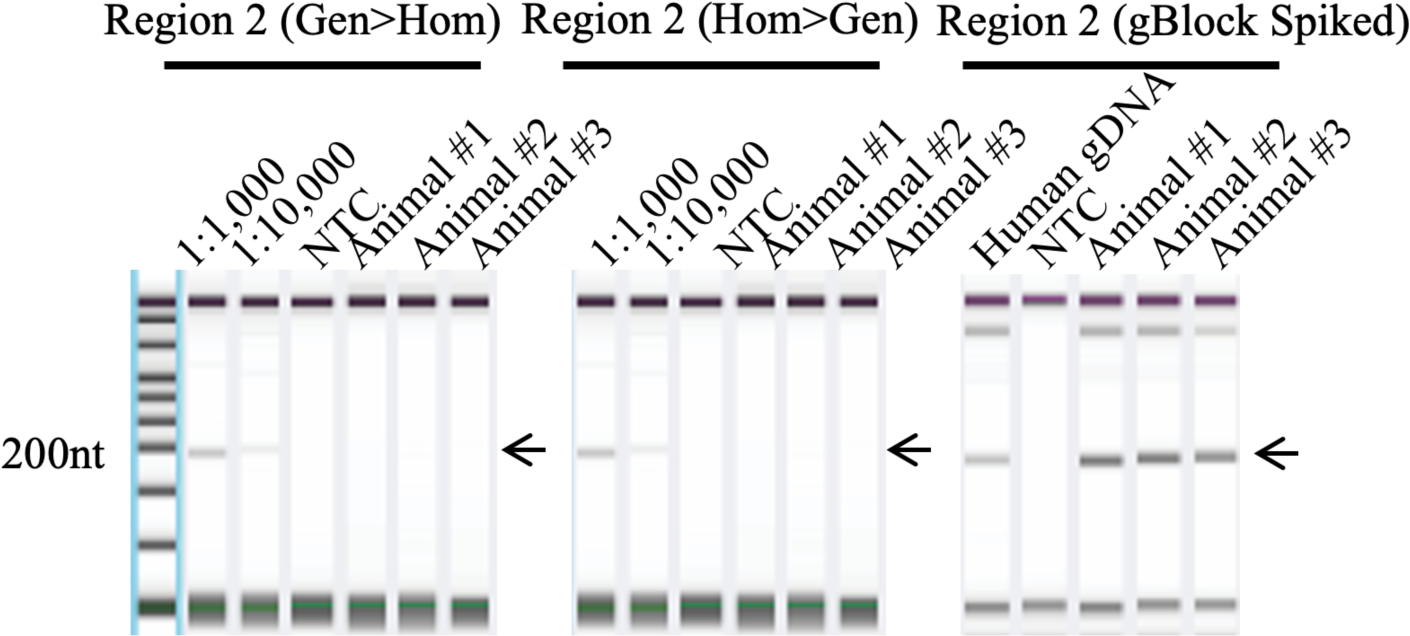
Analysis of Predicted Off-Target Integration. Human genomic DNA was amplified for five predicted off-target sites (Supplemental Table 1). TapeStation profiles for Region 2 are shown. Positive control gBlock DNAs are shown for spikes of 1:1000 and 1:10,000 with PCRs from the genomic region (Gen) to the right of the potential integration towards the homologous region (Hom) and from the homologous region to the genomic region to the left side of the possible integration. Genomic DNA from three animals treated with vector is shown for these amplifications and for a No Template Control (NTC). In the right panel, human genomic DNA and DNA from the same three animals are amplified in the presence of the positive control gBlock DNA to demonstrate that amplification would be possible if the off-target integration had occurred.

### Genome-wide Integration Assay (GWIA)

#### Amplification specificity

Examination of predicted editing sites is the only measurement done when some editing technologies are evaluated.^27^ However, it is preferable to examine all potential integration sites more broadly to characterize the editing/integration process fully. When scanning the entire genome for any potential integration sites, selectivity can be achieved by using a specific primer for the vector payload, but the same degree of selectivity is not possible for the unknown genomic sequence adjacent to uncharacterized off-target integrations. Furthermore, the payload-specific primers provide specificity relative to genomic DNA, but episomal DNA contains the identical sequences as the integrated DNA so that primer provides no specificity relative to the problematic contaminating episomes. These can be present at up to thousands of fold excess relative to integrated DNA, creating a serious signal to noise issue. There are many methods to select for or against specific sequences (such as affinity tags), but we have not found these to be robust when used to select for or against AAV episomal DNA containing ITRs. Two techniques were found to aid specificity in this situation, the use of splinter adaptors attached to the genomic end of the DNA and use of chimeric Locked Nucleic Acid (LNA) oligonucleotides to selectively block episomal amplification at episome-specific regions. Both methods have been used previously during the characterization of other viral integrations.^28^

While the payload primer on one end of the amplified integration junction provides specificity for that DNA end, the primer on the other end must be sequence-agnostic to allow the detection of any region in the genome. This specificity limitation can be partially ameliorated via the use of splinter adaptors.^28^ Splinter adaptors are a pair of oligonucleotides that are partially complementary to each other. They consist of a longer DNA with an extended 5’ end and a shorter DNA that is blocked at its 3’ end to prevent extension even in the presence of the longer sequence. These adaptors are ligated to the repaired ends of all free DNAs. Amplification starting from the splinter end of the DNA can only be achieved if the 5’ extension is initially replicated from another priming site located nearby (schematic in Supplemental Figure 2). Ideally, activating replication occurs only when the payload-specific primer is extended, but other artifactual sources of copying could also happen and replicate the longer splinter adaptor, making it available for artifactual priming. Nonetheless, the payload-primer specificity at the vector DNA end is assisted by some degree of specificity at the genomic end of the amplicon using splinter adaptors.

Another method for further improving specificity is using LNA oligonucleotides to block the amplification of unwanted episomal DNA.^29^ Because LNAs bind tightly to DNA, they can prevent DNA amplification when 3’ blocked and bound to specific sequences. In each of our rAAV vectors, short sequences are located between the homology arms and the ITRs that are present in the episomal DNA but not in the HR-integrated DNA. Binding LNAs to this region does not affect amplification of HR-integrated DNA but significantly inhibits amplification of the episomal DNA. LNAs of 20-24 nt were found to be specific and effective for this purpose.^29^ Various other methods to improve selectivity were tested, but the challenging nature of rAAV was found to limit their usefulness for either positive or negative selection.

Using nested PCR with splinter primers and LNA blocking, the GWIA method (Figure 3) was developed. DNA containing the codon-optimized vector payload (CO-hPAH) fused to human genomic sequences irrespective of where those human genomic sequences are located is amplified preferentially and independently of location, so it is possible to simultaneously detect low levels of both on- and off-target integration. Unfortunately, like other integration assays, GWIA is subject to artifacts so appropriate controls need to be included to avoid problem sequences like telomeric DNA that may self-prime, mitochondrial DNA that is present at thousands of copies per hepatocyte^30^ and repeat sequences that may misprime off each other. Additionally, unintegrated episomal VGs can significantly disrupt amplification due to their appetite for vector-specific primers. The excess vector DNA also contributes to two primary sources of artifactual signals that interfere with the detection of integration events. First, the episomal DNA can ligate to fragmented genomic DNA during sample processing, generating a false positive integration signal. Second, cells containing high levels of VGs can create a multitude of concatemers and other recombinants in which VG DNA may be rearranged into configurations that allow efficient amplification with one or two primers, increasing noise, reducing primer availability, and making true positive signals hard to distinguish. The degree to which these artifacts obscure the detection of true signals depends on the nature and concentrations of VGs and must be evaluated in multiple conditions so that the effective detection range of GWIA can be determined.

**Figure 3:**
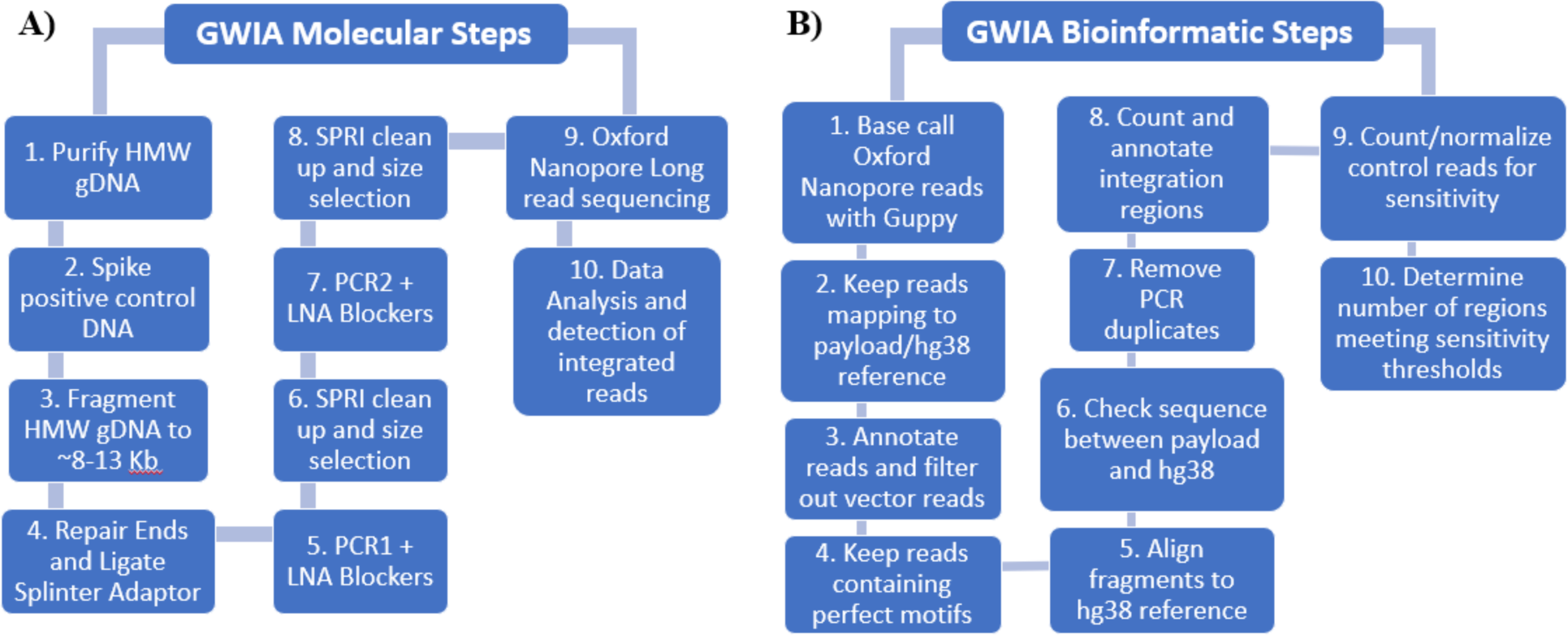
GWIA Schematic. Schematic flows for laboratory and informatics steps in the GWIA methods are listed. A more complete listing of details is provided in methods.

### Positive control DNAs

On-target integration occurs at up to ∼6%, but off-target integration, if it occurs at all, is much less common, making it more challenging to detect. Thus, generating a positive control for determining the LoD is imperative. There are multiple literature examples of purported viral integration findings that were later challenged by others as being caused by artifacts rather than real integrations,^15,31,32^ and it would not be surprising to find additional examples with AAV, especially with samples rich in vector episomes. Sample and library processing can generate various artifacts that share some properties with integration events but are not authentic integrations. To address this, we have developed two types of positive control DNAs for assessing LoD: a series of plasmids containing pseudo-off-target integration junctions and a cell line with an on-target integration that includes SNVs to distinguish it from authentic on-target integrations. For the plasmid positive control off-target DNAs, 36 random fragments of human genomic DNA were cloned adjacent to the right homology arm. These plasmids were sequenced to identify the origin of their genomic DNA, and they were found to arise from diverse genomic regions (Supplementary Table 2).

While the control plasmids allow us to query many different sequence contexts, independently prepared, high quality plasmid DNA may behave differently than the longer genomic DNA. To assess sensitivity and specificity using genomic DNA, a cell line was generated to provide an additional positive control for on-target, genomically integrated vector. A plasmid containing the payload region with variants to distinguish it from the standard gene therapy vector was co-transfected with CRISPR-Cas9 molecules containing sgRNAs targeting two sites in the chr12 region at which integration was intended. In a fraction of the cells, CRISPR-Cas9 cleaves the targeted locations which can then be repaired using the plasmid with the slightly variant homology arms. Because this repair could occur via Non-Homologous End Joining (NHEJ) and may not be HR-directed, this recombination may be different than what would occur with the virus and errors might be introduced in or near the integration site. Other off-target, CRISPR-dependent sites might also generate integration events.

After transfection of the plasmid and CRISPR-Cas9, cells were single-cell cloned and expanded. The resulting individual clones were screened by PCR for the desired chr12 integration event. Clones that appeared to contain the correct junction DNA by PCR were sequenced using Oxford Nanopore. One cell line with an appropriate integration (A11) was found which also contained normal genomic DNA in the same region, indicating the integration event had occurred in only a single chr12 allele. In addition to this planned integration event, another complex event was detected via GWIA that included a segment of chr17 and multiple vector payloads. Whole-genome sequencing of A11 with PacBio was performed to characterize the integration pattern better. In addition to the expected CRISPR-driven integration event on chr12, the second, unexpected integration event included >2 kb from chr17 and multiple payloads and vector cloning sites but only a small segment of ITR. The complexity and length of this integration region prevented complete assembly. DNA prepared from this cell line has been used as a positive control by spiking into assays of test genomic DNA at known amounts to evaluate the sensitivity of the GWIA method for detecting integration.

To confirm the blocking ability of LNAs in this system, genomic DNA from A11 was amplified using three primers. One primer is in the PAH payload sequence and is present in both the on- and off-target integrated DNA. The other primers were designed to match adjacent genomic DNA from the intended on-target site in chr12 and the observed off-target site in chr17. The chr17 off-target site includes a vector sequence not present in the HR-like integration event on chr 12 and the blocking LNAs target this vector sequence. When no LNAs are present, these DNA fragments are amplified equally efficiently whether alone or together (Supplemental Figure 3, lanes 1-3). When the vector-specific blocker is added (lanes 4-6), chr17 amplification is blocked whether alone (lane 4) or in combination with the chr12 primer (lane 6), confirming that LNA can specifically block vector sequence amplification.

### Limit of Detection (LoD)

The 36 different plasmid control DNAs with artificial integration junctions were spiked into A11 genomic DNA at varying relative concentrations and subjected to GWIA to assess both on-target integration provided by the A11 DNA and off-target integration provided by the 36 different plasmids. GWIA is described in detail in Materials and Methods with a schematic of the laboratory and analysis workflow shown in Figure 3. Briefly, genomic DNA is spiked with appropriate controls, sheared to ∼8 kb, repaired, ligated to splinter adaptors, amplified with one pair of payload/splinter primers in the presence of LNA blockers, size selected, amplified a second time with a different, nested pair of payload/splinter primers in the presence of LNA blockers, size selected, and sequenced using Oxford Nanopore flow cells. The resulting data is then base-called, aligned to vector and hg38 references, and filtered based on multiple sequence motifs.

When plasmids are spiked into A11 positive control genomic DNA and analyzed relative to on-target reads from A11, the recovery of plasmid “integration” reads is consistent across all sequence types (Figure 4). This shows that it is possible to detect off-target integration events mimicked by the plasmids across a wide variety of sequence contexts when those integration junctions are present. To further assess the assay’s sensitivity for low levels of integration junctions, varying concentrations of the different plasmids were spiked into genomic DNA over a 1000-fold range starting at a 1% molar ratio relative to genomic DNA and going down to 0.001%. Detection over a 100-fold range was reasonably linear (Figure 5), but the lowest concentration was not readily detectable in these conditions. Thus, when the DNAs are of good quality and unencumbered by contaminating VGs, detection of integration junctions down to 0.01% can be readily achieved.

**Figure 4:**
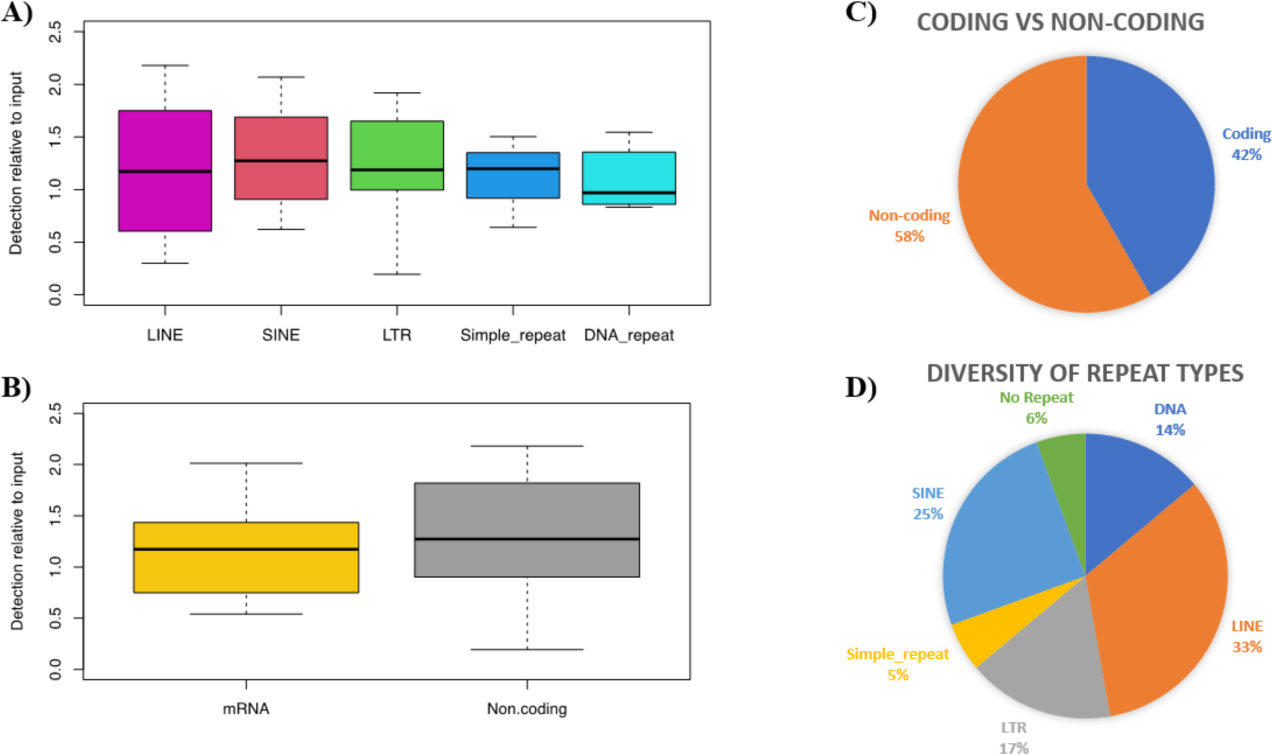
Read Frequency for Positive Control Plasmids using GWIA. Sequences cloned adjacent to RHA were classified as their repeat content and whether they contained coding or non-coding sequence. Relative read counts for each junction were made and averaged by class.

**Figure 5.**
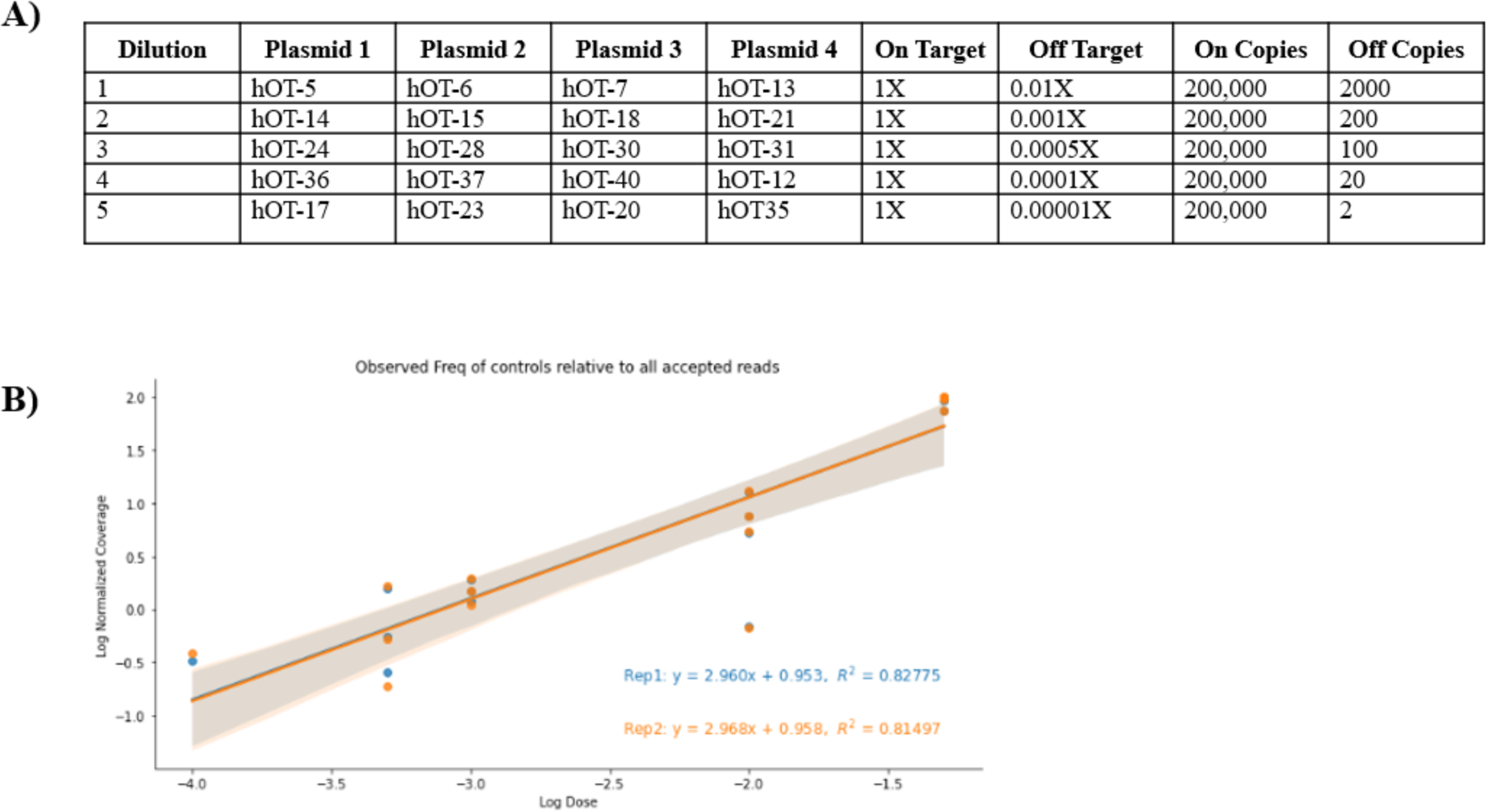
Detection of Control Plasmid DNAs. Twenty plasmids with RHA-genomic fusions were split into four groups and diluted as listed in A. Each mix was sequenced and reads were assigned to the different plasmids. The correlation of dilution and read count is shown (B).

### Limit of Blank (LoB) in the presence of VGs

AAV capsids lysed via heat treatment release both plus and minus strand ssDNA and the complementary strands rapidly anneal to generate dsDNA from the +/− ssDNAs. This dsDNA can be spiked into reactions to assess VG-dependent artifacts. However, because the VG DNA was never in cells after packaging, the spiked DNA does not contain any integrated or concatemeric DNA like that potentially generated inside cells. Control genomic DNAs from either human or mouse were spiked with varying concentrations of lysed vectors and A11 genomic DNA (Table 1) to mimic the composition of DNA in transduced cells with no non-control integrated DNAs present. The lysed vectors were spiked before fragmentation, but their smaller starting size relative to genomic DNA makes it likely they are minimally affected by the shearing.^33^ In these samples, the starting molar ratio of vector to human DNA ranges from 100:1 to 10,000:1. Depending on the injected vector dose, the actual ratio in hepatocytes treated with high doses of rAAV can be up to 1000:1. With the highest ratios of vector DNA, very little human DNA is detected because most of the reads are episomal DNA that likely undergoes linear or inefficient amplification. As the amount of lysed vector is decreased, the number of RHA reads increases as well as the number of reads with both RHA and hg38 sequence. The amount of linearly amplified vector decreases while the human DNA is amplified better. However, even with the lowest amount of lysed vector, nearly 90% of reads contain neither RHA nor other human sequences, showing the preponderance of artifactual vector sequences that are amplified despite efforts to minimize their impact.

**Table 1:**
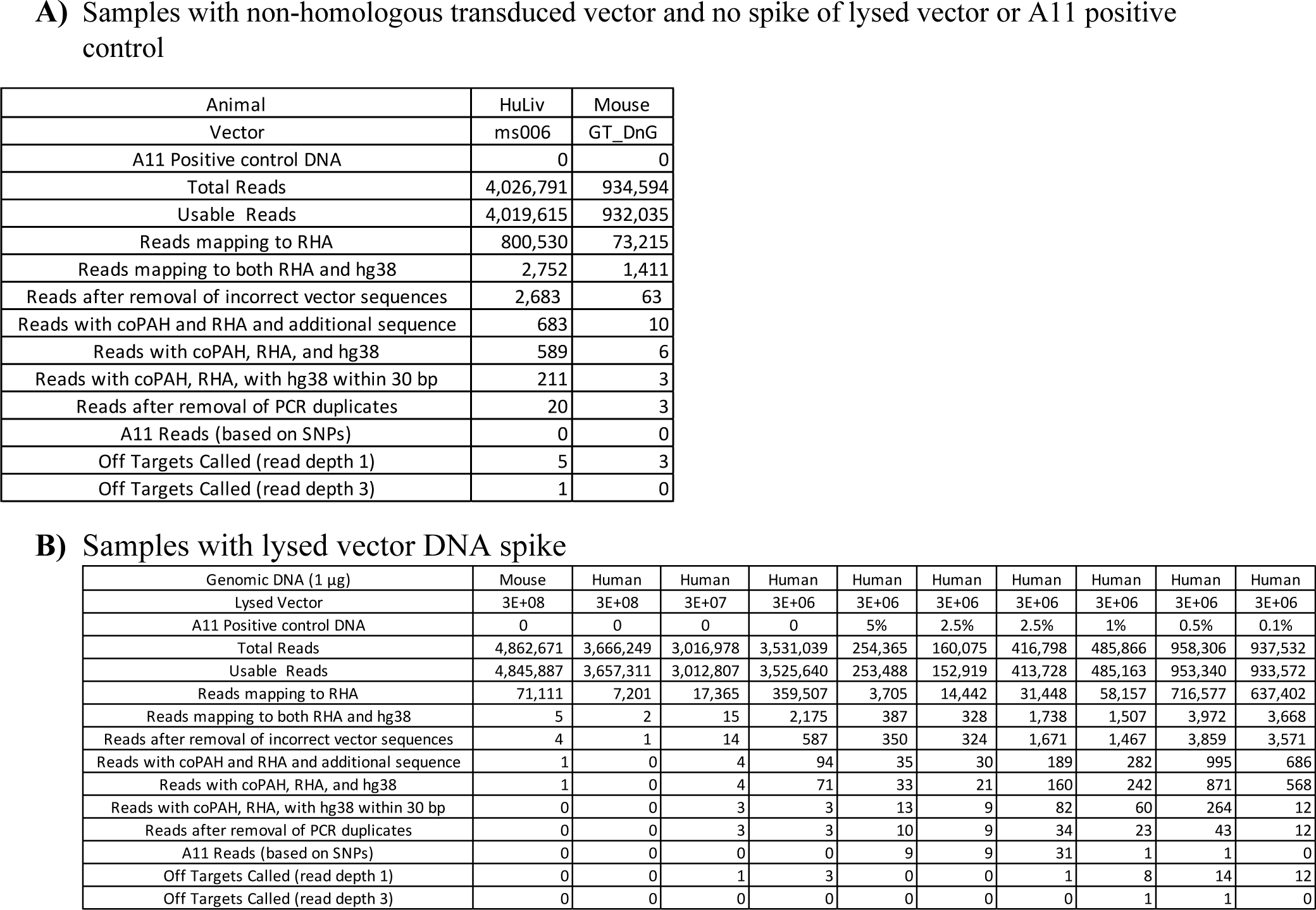
Limit of Blank using control DNA spikes in control genomic DNA.

When A11 genomic DNA is also added along with the lysed vector, the number of total reads drops substantially, but, depending on the amount of A11 added, the number of reads corresponding to RHA and hg38 increases. The number of reads with CO-PAH, RHA, and hg38 sequences in the correct locations for integration is much higher with the A11 spike. However, amplification of the A11 integration site is clearly non-linear as very few A11 reads are seen with 1% A11 DNA added while more than 10x as many reads are seen with 2.5% and 5% A11 DNA. Some sequences also correspond to off-target sites, but almost all of them are singletons that are not conserved across samples. Because no integrated DNA can be present, these must all be false positives arising from artifactual processes. The presence of false positive integration signals in DNA controls where no integration could have occurred highlights the need to determine the LoB in different conditions. Furthermore, any suspected off-target integration event should be confirmed in tested samples via PCR with custom designed primers.

### GWIA in Engrafted Human Hepatocytes

The use of control plasmid and genomic DNAs to model the behavior of DNA from treated cells could be incomplete because the rAAV DNA from capsids transduced into cells can undergo complex rearrangements that are not readily mimicked by the control DNAs prepared in vitro. To better understand the impact of in vivo VGs, A11 control genomic DNA was spiked into genomic DNA isolated from human hepatocytes from engrafted FRG mice treated with varying amounts of vector for different times post-injection. These samples contain the full range of DNA species including episomal vectors, concatemers, and other vector rearrangements that may arise over time in cells. The various DNAs may behave unexpectedly during library preparation, leading to artifacts including non-specific ligation to other DNA fragments and primer-independent (via ITRs) or single primer amplifications. As shown in Figure 6, increasing concentrations of A11 control DNA spiked into the samples from mice dosed at 7E+13/kg yields a reasonably linear response. The episomal VG load in these cells was determined using qPCR measurements. This assay does not distinguish between monomeric episomes and more complex concatemeric structures. The low-dose samples average 47 CO-PAH copies /allele. At the higher dose, 2E+14/kg (average of 79 CO-PAH copies/allele), the response is no longer linear and flattens above 3% A11 DNA. Thus, it is possible to detect the integration junctions but not accurately estimate their frequency using GWIA in high VG conditions.

**Figure 6:**
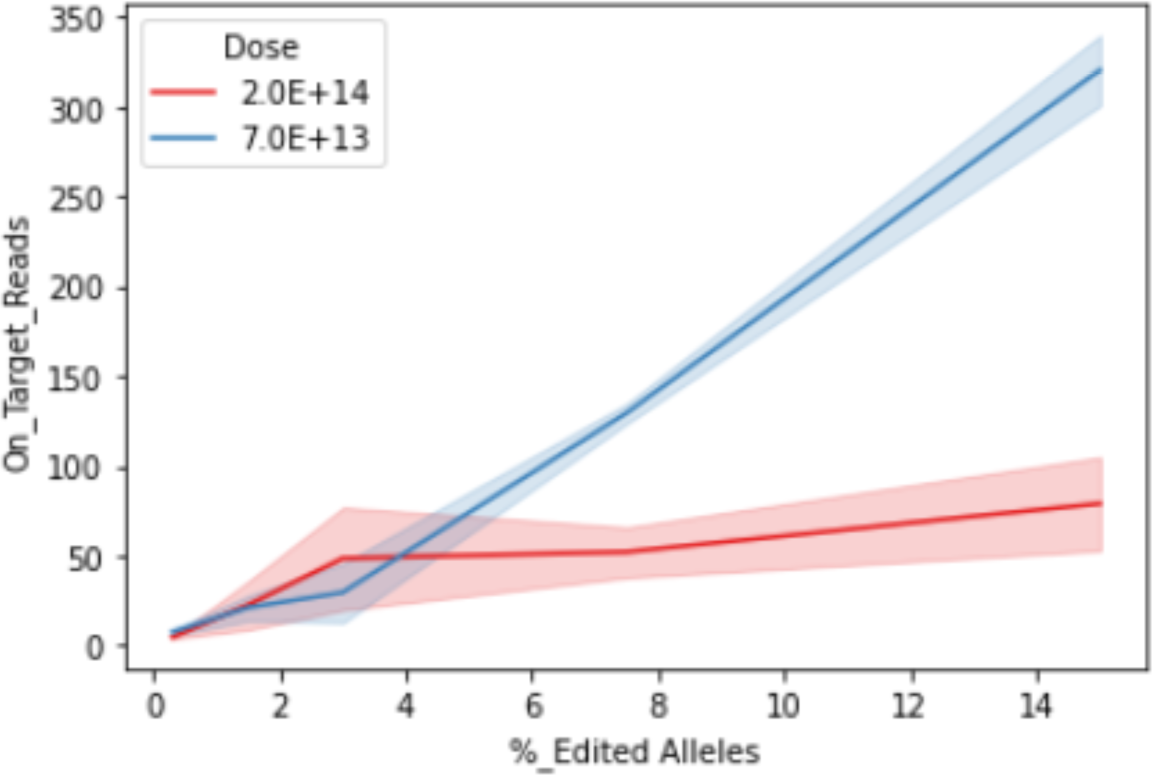
Limit of Detection in FRG Mice with A11 Spike. On-target integration reads from A11 were counted for all FRG mice treated with hHA-LP1-hPAH. The number of A11 reads as a function of spiked DNA fraction is shown for the low- and high-dose animals.

It is also instructive to examine reads that are filtered out during informatic analyses. By far, the two filters that eliminate most of the reads require that the resulting sequences have regions that map to the RHA and a separate region that maps elsewhere in hg38. This eliminates over 99% of reads in all cases and highlights the rarity of events sought. Other filters that have a lower impact include those eliminating reads with vector sequence beyond RHA and those where the RHA and other hg38 sequences are separated by more than 30 bp. When the intervening sequence is longer than 3 bp, it is usually random and cannot be aligned. Authentic junction sites tend to have 0-3 bp between the RHA and hg38 regions with that variable segment generally caused by sequencing errors. PCR duplicates, which are assigned based on the positions of the RHA and gDNA breakpoints, are also removed, but these are infrequent given the low number of junctions found.

### Off-target integration in engrafted hepatocytes

Based on the findings that 2.5% A11 DNA spiked into genomic DNA from engrafted human hepatocytes would provide the best sensitivity in the treatment range we performed, we examined genomic DNA purified from human hepatocytes from mice four or twelve weeks after injection. FB-treated mice had no episomal VGs and thus could not generate concatemers that could amplify and compete with the integration signal. FB-treated mice had ∼100x more positive control reads than hHA-LP1-hPAH treated mice. Additionally, the low-dose hHA-LP1-hPAH treated mice had more positive control reads than the high-dose hHA-LP1-hPAH treated mice due to lower interfering VG levels. All replicate samples had on-target reads that were within 2x of each other. Only the on-target integration site on chr12 generated more reads than the positive control cell line genomic DNA used to establish LoD. No genomic regions had reads higher than the 0.5% LoD based on the number of reads from the spiked genomic DNA. The location of integration junction reads is shown in Figure 7. While the intended site is easily observed, all other reads were sporadic and likely due to library artifacts. If any significant off-target sites had been identified, PCR primers would have been designed to determine whether the purported events were real.

**Figure 7:**
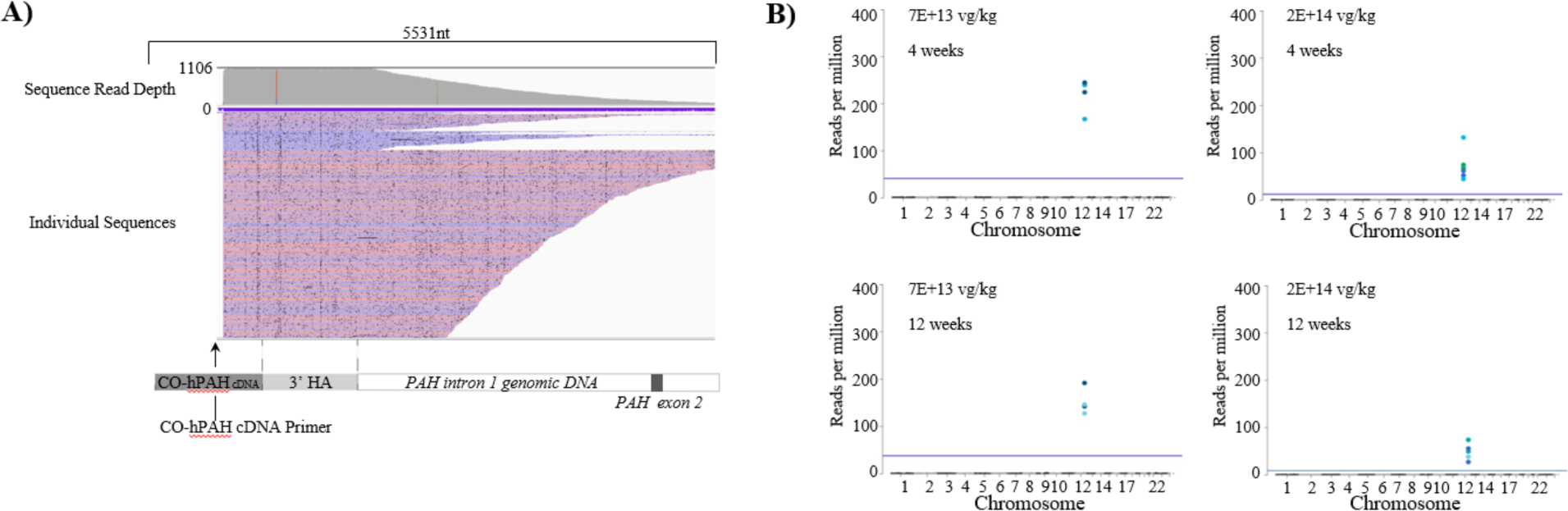
GWIA in Transduced Engrafted Human Hepatocytes. A) IGV view of reads mapping to the on-target integration site on chr12. Each line corresponds to an individual, non-duplicated read. Blue and pink colors indicate the orientation (+/−) of reads. The map of the integration region is shown below the reads. B) The on- and off-target reads for each animal are shown in a Manhattan plot. The LoD is shown as a horizontal line. Only the on-target reads are visible because off-target reads are too infrequent.

The diversity of sequence ends is shown in a representative Integrative Genomics Viewer (IGV) file displaying ONT sequencing reads from one low-dose hHA-LP1-hPAH sample (Figure 7). Each single molecule read contained sequence starting with the CO-hPCR primer located on the left end and going through the RHA region into adjacent genomic DNA (up to thousands of bases depending on the position of the shearing site) and ending with the adaptor ligated to the end of the randomly fragmented genomic DNA. The range in length distributions observed on the right end of the reads is caused by the reverse primer binding to the adaptor ligated to the randomly sheared ends of different molecules.

Many artifacts arise in an unknown manner and could come from events that occur in the cell or during subsequent library preparation, ligations, and amplifications. Three reads from the FRG mouse samples are interesting because they are fusions of lambda DNA with human DNA. If lambda had been injected along with rAAV, these might be ascribed to lambda integration into human genomic DNA. However, lambda is only present at the last stage of library preparation where it is added as an ONT sequencing control and is found well below the LoD. These findings highlight the potential for ligation or PCR-induced artifacts arising during sample processing.

## Discussion

AAVs have shown durable expression in the liver using AAV gene therapy for hemophilia, with human data out to 7 years^34^ and canine data out to 10 years.^35^ More recently, AAV delivery of a functional *PAH* gene has been shown to restore the normal biochemical pathway in a mouse model of PKU and is currently in clinical trials for adults with classical PKU.^24^ The preclinical studies demonstrated durable expression of human PAH in PAH^enu2^ mice and normalization of Phe out to a year post-dose (end of study). This approach used AAV-based gene therapy to deliver and express the gene extra-chromosomally in episomes. As the transduced cells divide, dilution of the episomes can occur given that the episomes do not replicate with cell division. While hepatocytes in the adult liver divide over time, this process is relatively slow compared to a growing liver during childhood development. For a disease like PKU that is diagnosed shortly after birth, a gene editing-based therapy with durability through liver growth and hepatocyte turnover may benefit adults as well as younger individuals.

Several approaches aiming to correct the underlying mutation in PKU have been reported. Villiger et al^36^ showed that using two AAV vectors, delivery of a Crispr-Cas9-base editor targeting the PAH^enu2^ mutation could correct the mutation in the liver resulting in a reduction of serum Phe to normal levels. Richards et al^37^ showed that using a dual vector approach delivering two doses (at 1 and 5 weeks of age) of a Crispr-Cas9 nuclease specific to exon 7 downstream of the PAH^enu2^ mutation and a 2.7 kb repair template encoding the corrected variant resulted in serum Phe lowering that was sustained for 65 weeks. This approach relied on the homology-directed repair (HDR) mechanism to correct the PAH^enu2^ mutation. In this case, the percentage of HDR-modified alleles averaged 13%, with 7% showing functional correction at the cDNA level, and resulted in Phe reduction, but not to normal levels. More recently, Bock et al^38^ showed that using AAV8 and AdV to deliver a size-reduced *SpCas9* prime editor could achieve 11% editing in the PKU model, resulting in serum Phe concentrations below the therapeutic threshold of 360 µM.

While these approaches are promising, they target a specific mutation in the *PAH* gene and are not directly applicable to the thousands of other *PAH* mutations. Furthermore, like many other CRISPR-based approaches, potential off-target events were not examined in a whole-genome manner. Others have easily identified large structural effects induced by CRISPR.^39^ However, potential single base changes cannot be readily evaluated across the whole genome because the background level of single base changes that occur with every round of replication can obscure the signal from such potentially deleterious modifications.^40^ Targeted assessment of the most likely hot spots for editing can be carried out, but most of the genome remains unevaluated for mutations. HR-driven AAV integration events are easier to detect and distinguish from any background activity because of their larger size and unique sequence characteristics.

Two particularly important artifacts must be addressed when attempting to identify real AAV or other viral integration events: 1) Processing-dependent ligation of AAV fragments to other DNAs that may appear to be fusion events but are simply in vitro ligation artifacts; and 2) use of short read sequencing technologies to assemble long molecules that may not actually be present when examined as long reads. Indeed, a study comparing NGS methods that included controls showed that ligation was a significant driver of false positives in control samples.^19^ Use of transposase instead of ligase reduced artifacts.

Because of the high background of AAV genomes and their propensity for generating concatemers, amplification of minor components can lead to unexpected fragments whose source could be attributed to integration rather than be correctly identified as artifacts. Some artifacts, like PCR mispriming, can be readily detected by appropriately examining the sequenced DNA^23^ while others can be more challenging. Because our nested PCR primers do not extend to the end of the splinter adaptors, we can eliminate mispriming as a significant cause of artifacts in our system because those sequences are not present in our products. We have included relevant positive controls and long-read sequencing in our methods to ensure that we can distinguish real integration events from artifacts at our stated sensitivity.

Awareness of artifacts in the study of viral integration is recognized as an issue in the literature. Kaeppel *et al.* (2013)^41^ reported a significant number of mitochondrial-AAV integrations, but this result was challenged by Cogne et al (2014),^15^ who used an alternative approach to minimize artifacts. The response to this challenge cited new experiments with controls intended to address the issues raised, but the new controls either contained no AAV DNAs or no primers that would allow their amplification. Thus, the original issues raised remain relevant and it is possible that the insertion events identified could be simply processing artifacts and not actual integration events. Similar integration artifacts have been observed and challenged when looking at other putative viral “integrations”.^31,32^ Any potential off-target integration events need to be confirmed using an orthogonal method, like directed PCR, before citing their occurrence.

While it would be desirable to achieve much greater sensitivity when examining off-target integration, it is not generally possible to do so with the limitations of sample availability and the sequence similarity with episomes. While efforts to improve sensitivity will continue, it is worth noting the safety record of rAAV across systems. After 10 years post-administration of AAV8 or AAV9 vectors expressing canine factor VIII, none of the dogs showed evidence of tumors, altered liver function, or any liver pathology features that could be attributed to treatment with AAV.^42^ George et al. published a follow-up study in humans treated with AAV expressing Factor 9 and also found no evidence of liver toxicity or other major safety concerns out to 15 years post-dose.^35^ An earlier study concluded the same lack of hepatic genotoxicity in nonhuman primates and humans.^43^

A hallmark of AAV-mediated HR is the precision of integration and avoidance of unwanted mutations. HR is the preferred DNA repair pathway for the gain-of-function insertion of DNA given the precision and lack of insertions/deletions that could disrupt expression as seen with nucleases and base editors. Previous work using short-read NGS showed that AAVHSC15-mediated integration did not introduce indels or ITR sequences at the target integration site.^22^ The use of long-read sequencing is critical in these studies to ensure that genomic DNA is being analyzed and not the much higher copy number episomal DNA. Confirming the desired integration is a crucial safety assessment for integrating vectors. When evaluating across the genome using long-read NGS, the only site of DNA integration detected above the LoD was at the desired human *PAH* locus.

## Methods

### Animal Procedures for Mice

Housing, breeding, and all procedures performed on PAH^enu^^2^ mice were in accordance with the Institutional Animal Care and Use Committee (IACUC) at Homology Medicines, Inc.

The FRG mice were housed in an IACUC accredited facility. General animal care and housing procedures are described in Guide for the Care and Use of Laboratory Animals, National Research Council, 2011, Yecuris™ IACUC Policy, and Yecuris™ General Mouse Handling Care and Euthanasia. Cage changes occurred every 2 weeks and the testing facility was sanitized weekly. Animals were provided irradiated mouse chow with a low Tyr content (0.53%, PicoLab® High Energy Mouse Diet, 5LJ5 chow) and drinking water ad libitum. Animals were administered the prophylactic antibiotic sulfamethoxazole and trimethoprim (SMX/TMP, 80 mg/mL/16mg/mL) every other week in their drinking water. They were administered 2-(2-Nitro-4-trifuoromethylbenzoyl)-1, 3-cyclohexanedione (Nitisinone, NTBC) for 3 days once every month.

### Fah*^-/-^*/Rag2*^-/-^*/Il2rg*^-/-^* (FRG®) Humanized-Liver Xenograft Mouse Model

The Fah^-/-^/Rag2^-/-^/Il2rg^-/-^ (FRG®) mouse model has been characterized by Azuma et al.^26^ In brief, engrafted human hepatocytes integrate completely into the structure of the recipient mouse liver and occupy greater than 80% of the parenchyma without disturbing the recipient liver organization. Human hepatocytes can be distinguished morphologically from mouse hepatocytes. From a functional perspective, engrafted human hepatocytes retain the functional properties of mature, differentiated hepatocytes as demonstrated by the expression of mature hepatocyte-specific genes and albumin secretion. Typically, 75% of the transplanted mice achieve and maintain high levels of long-term engraftment, demonstrated by histology, DNA analysis and enzyme assays. The majority of the mice that survive the procedure have high levels of engraftment based on circulating human albumin levels and post-mortem histology.^44^

To generate humanized livers in FRG mice, human primary hepatocytes are implanted concurrently with the withdrawal of 2-(2-Nitro-4-trifuoromethylbenzoyl)-1, 3-cyclohexanedione (Nitisinone, (NTBC) that is required for survival by mouse hepatocytes in this strain. Since the human hepatocytes have an intact fumarylacetoacetate hydrolase (FAH) gene, only mouse hepatocytes are affected by the loss of NTBC, resulting in the gradual repopulation of the mouse liver compartment with human hepatocytes.^26^

FRG mice achieve high levels of repopulation 5-6 months post-transplant and are typically used within 6-8 weeks to avoid natural loss of engraftment or tumor development as they age. Compared to wild-type mice, the FRG mouse model lacks B-cells, T-cells and NK-cells due to the absence of the IL2rg and Rag2 genes. In addition, the remaining mouse hepatocytes do not metabolize Tyr due to the lack of the *Fahah* gene. Despite these changes, this mouse model has proven valuable in many research areas including infectious diseases, NASH, gene therapy, metabolism and toxicology.^45–48^

The in-life and hepatocyte isolation portion of this study was conducted by Yecuris (Tualatin, Oregon) who generated this model and have specialized technical expertise in handling this mouse. Due to the complexity of the generation and handling of this model, the number of FRG mice that can be utilized in a single experiment is limited.

#### Injections

4- and 10-week-old PAH^enu2^ mice were weighed prior to injection, and a single I.V. injection of the test article was administered retro-orbitally for all studies. Mice were anesthetized using isoflurane for the injection. Proparacaine was applied to the eye before injection. The needle was inserted into the retro-orbital sinus via the medial canthus and the test article was injected slowly into the sinus. All doses were administered at 10 mL/kg.

FRG mice were removed from NTBC for ≥ 25 days and SMX/TMP for ≥ 3 days before dosing to reduce the number of mouse hepatocytes. Mice were weighed before dosing and a single I.V. injection of the test article was administered retro-orbitally using an insulin syringe (VWR, Radnor, PA). One day post-dosing, the mice were put on NTBC and continued with the standard NTBC water cycle for the duration of the study.

### Design and construction of AAV vectors

The same AAV backbone plasmid, which included 5’ and 3’ AAV ITRs and flanking *PAH*-targeted integration sequences, was used for all vectors. These vectors include a CO-hPAH DNA flanked by locus-specific arms identical to human genomic sequences. The human 5’ homology arm aligns to human genome build hg38, chr12: 102916857-102917816 and the 3’ homology arm aligns to hg38 chr12: 102915806-102916716. The homology arms were selected to be unique in the genome with no sequences of 40 nt or longer having homology to other locations. A late SV40 polyadenylation sequence was included as a transcription termination sequence. Promoter-containing vectors include the liver-specific promoter LP-1 as previously described,^49^ positioned between the 5’ homology arm and the CO-hPAH DNA. The LP-1 promoter, polyadenylation cassette and CO-hPAH DNAs were synthesized as gBlocks by IDT. The ITR containing plasmid backbone, the CO-hPAH DNA, homology arms and regulatory sequences were assembled by standard restriction digestion and ligation techniques.

### Test articles

All vectors used were manufactured at Homology Medicines via a transient transfection process using a HEK293 cell line. The vectors were packaged in AAVHSC15 capsid and are comprised of an expression cassette as described above with or without the LP-1 liver-specific promoter, human homology arms, and AAV2 ITRs on both ends of the construct. Vectors were diluted in a pH-neutral buffer for I.V. administration. Vectors were tested for endotoxin (<10 EU/mL) and titered by Droplet Digital™ polymerase chain reaction (ddPCR) using primers targeting the CO-hPAH DNA. Vectors were analyzed for VP1, 2, and 3 ratios by silver- and Coomassie Blue-stained SDS-PAGE and capsid titer by ELISA.

### Vector genome copy number by qPCR

A real-time quantitative polymerase chain reaction (qPCR) method was used to quantify vector genomes in mouse liver samples or isolated hepatocytes from FRG mice. Samples containing gDNA and control plasmids were prepared for analysis on the ThermoFisher QuantStudio™ Flex Real-Time PCR System, which utilizes TaqMan technology. The assay’s linear range was 50 to 10E+8 copies per reaction with an LLOQ of 50 copies of vector per 1 µg of gDNA. Samples were quantified against linearized plasmid. Real-time PCRs were performed on 96-well plates, and each plate was run with a standard curve with no template control and study samples. The standard curve was run with the plasmid in 3 replicates for the standard points at 1E+9, 1E+8, 1E+7, 1E+6, 1E+5, 1E+4, 1E+3, 1E+2, 10 and 1 copy per reaction. Vector-specific sequences were amplified by real-time TaqMan PCR using the primers and probes for CO-hPAH (extended methods). The PAH probe consists of the FAM™ fluorescence reporter dye at the 5’ end of the probe, the internal ZEN™ quencher (between the 9th and 10th nucleotides), and the Iowa Black® fluorescent quencher at the 3’ end of the probe.

### Analysis of on-target integration frequency by Oxford Nanopore sequencing

On-target integration frequency was determined using a primer competition assay (3-primer). The relative frequency of integration and non-integration sequences were tallied by long-read Oxford Nanopore sequencing. The raw integration frequency was calculated as the percentage of integrated reads divided by the sum of integrated and non-integrated sequences. The raw frequency was corrected for a length-dependent PCR efficiency adjustment using independently amplified DNAs and reduced by an additional factor caused by the amplification of concatemeric sequences that interfered with the amplification of non-integrated sequences..

Each reaction included 100 ng of input genomic DNA. The frequency of integration by PCR was normalized to a PCR efficiency curve. DNAs representing wild-type and integrated DNA were mixed in known ratios. These templates were diluted and assayed by the PCR reaction as described.

Purified WT and edited control templates were quantified with the Qubit 1X dsDNA HS assay kit. The linear detection range of this assay kit is 0.1 ng/μL to 100 ng/μL. A stock control dilution was made for WT and edited control templates in water, 1 ng/μL in 100 μL volume.

A concentration of 0.1 pg/μL was used as a working dilution for testing amplicon efficiency. 0.1 pg/μL contains ∼70,000 copies of control template molecules. The amplicon efficiency test was set up with 50% WT and 50% edited control templates with 0.1 pg total (0.05 pg WT and 0.05 pg Edited), so that the molecular number for the WT template is ∼35,000 copies.

The resulting amplicons were prepared for Oxford Nanopore sequencing in accordance with the manufacturer’s instructions:

“Genomic DNA by Ligation (SQK-LSK109)” used for single-plexing: https://community.nanoporetech.com/attachments/3041/download
“Native barcoding genomic DNA (with EXP-NBD104, EXP-NBD114, and SQK-LSK109)” used for multiplexing with up to 24 barcodes: https://community.nanoporetech.com/attachments/3186/download

Each amplicon to be sequenced was cleaned using AMPureXP beads and samples were quantified using a Qubit 1X dsDNA HS kit. 200 fmol of input DNA was end-repaired. Each library was purified by AMPureXP magnetic bead cleanup per the manufacturer’s instructions. Up to 200 fmol of DNA was ligated to Nanopore barcoded adapter mix II and T4 DNA ligase followed by AMPure bead cleanup and quantified by Qubit. DNA was prepared for final loading with 37.5 µL of sequencing buffer 25.5 µL of loading buffer and 12 µL of DNA library per flow cell. Each library was loaded into the SpotON sample port and sequencing was initiated per the manufacturer’s instructions.

#### Analysis

The resulting Fast5 files were base-called using Guppy basecaller. Only reads of the appropriate amplicon size range between 1,100nt and 1,400nt were included. Reads which have plasmid or vector sequences such as ITR were removed. All remaining reads were competitively aligned to a WT and integrated genome reference sequence using the Minimap aligner. The raw read frequency of wild-type and integrated sequences were tallied. Calculated “raw observed integration %” = reads mapped to edited reference / (reads mapped to Edited reference + reads mapped to WT reference). The standard curve of (0%, 0.5%, 1%, 2%, 5%, and 10%) was used to generate linear regression formula for normalizing the raw observed % integration levels.

### Predicted Off Target

Each homology arm was broken into 80bp windows, sliding 10bp per window, for a total of 171 windows. Each 80bp window was queried against the human genome (Database: RefSeq Genome Database (refseq_genomes) Organism: Homo sapian (Taxid:9606)) with BLAST ((Expect threshold: 10, Word Size: 16, Match/Mismatch Scores: 1,−2, Gap Costs: Linear) to identify genome regions with >35bp match and >60% identity. From this analysis, 16 regions were chosen as primary hits. Primary hits were then queried again with BLAT (UCSC Genome Browser) to confirm the location and characterize the alignment. This narrowed the final list to 6 regions (supplemental table 1). One of the identified regions locates in a highly repetitive genome locus. We dropped this region out of the analysis because we couldn’t develop a reliable PCR based assay (region 3).

Off-target analysis was carried out with 10 ng of genomic DNA from three transduced Yucaris human hepatocyte samples. The amount of DNA available was limited so more could not be used. Each sample was amplified for 35 rounds with primers specific to the predicted inward and outward integrations for each identified region. Analysis of PCR products was done on a TapeStation to quantitate the anticipated bands. For each region, the relevant synthetic DNA positive control was spiked into the human DNA sample at two concentrations (1:1000 and 1:10,000) to ensure that the off-target band could have been detected if present.

### Genome-wide integration assay (GWIA) by long-read Oxford Nanopore NGS

GWIA experimental steps are summarized in Figure 3. gDNA from a positive control cell line containing the PAH edited allele with added SNPs was spiked into all samples at a 2.5% molar ratio. 500-1000 µg of total gDNA including spiked gDNA were randomly fragmented to ∼8-10kbp using COVARIS g-TUBES. Fragmented gDNA was end-repaired and A-tailed using the NEBNext Ultra II End Repair/dA-Tailing Module, and ligated to splinter adapter (GCCGAACATGAGCTATACGACTTAAACTTCCGCATGGCGTCTCCGCTTAAGGGACT; 5’Phos-GTCCCTTAAGCGGAG-3’AmMO) with NEBNext Ultra II Ligation Module.^28^ Ligated gDNA was first amplified with a payload-specific, biotinylated forward primer (5’-AAGACCGCCATCCAGAACTACA-3’) and an outer splinter reverse primer (5’-ACTTAAACTTCCGCATGGCGT-3’). LNA blockers (/5AmMC6/CTGCA+G+G+T+C+T+A+G+ATACGTAG/3AmMO/; /5AmMC6T/+AGATA+CGTA+GATA+AGTA+GCAT+GGC+G/3AmMO/; /5AmMC6T/+AGATA+CGTA+GATA+AGTA+GCAT+GGC+G/3AmMO/; /5AmMC6T/+AACCC+CTAG+TGAT+GGAG+TTGG+C/3AmMO/; /5AmMC6T/+GGCGT+CGGG+CGAC+CTTT+GG+T/3AmMO/) against non-payload VG sequences were used in PCR amplifications to limit episomal VG amplification. Biotinylated PCR1 products were size selected with 0.5X SPRI beads (SPRIselect) twice (to remove unwanted <500bp amplicons) and enriched by capture with streptavidin beads. Captured PCR1 products diluted at 1:100 were amplified again (nested PCR2) with the inner splinter primer (5’-CTTAAACTTCCGCATGGC GTC TC-3’) and an internal payload primer (5’-AGAGGATCGAGGTGCTGGATAA-3’). Finally, PCR2 products were size selected with 0.5X SPRI beads twice, and used to generate Oxford Nanopore libraries. Long-range sequencing was used to provide sufficient sequence coverage to confirm the payload CO-hPAH DNA, the complete homology arm, and adjacent genomic sequence. Sequencing data were analyzed with in-house developed computational methods tuned to filter out VGs and artifactual reads. The positive control cell line DNA spiked at 2.5% was used to estimate a lower limit of detection.

#### Analysis

After sequencing, reads were base-called with Guppy and quality filtered with Filtlong. Additional filtering was based on removal of the control lambda reads and any read that did not contain the vector payload or DNA matching human genome sequences. Sequences were annotated for components corresponding to vector. Reads with components that should only be present in the vector were eliminated. Reads supporting the possibility of being integration events were then filtered to eliminate sequences that did not meet criteria indicating they arose from legitimate integration events. Such reads must include all or some of the RHA as well as >200 nt following the 3’ end of the RHA. This non-RHA region is then aligned to hg38 and those with MAPQ<40 or length <100 nt are discarded. Reads with more than 30 nt separating RHA from hg38 are also discarded. PCR duplicates are collapsed to single reads. Control reads were identified and counted to determine positive integration read cut offs. Results from replicates were compared and only those appearing in both replicates were considered real with the others discarded as technical artifacts.

#### Data Availability

All fastq files used in this study have been uploaded to SRA under the accession PRJNA1001866.

## Notes

### Competing Interest Statement

All authors are current or former employees of Homology Medicines Inc.

### Summary of Updates

Correct misspelling of author name

