## supplemental tables and figures for "Genome-wide assays to characterize rAAV integration into human genomic DNA in vivo"

**[Title]**

### Supplemental Files

**Supplemental Figure 1:** Maps for vectors used

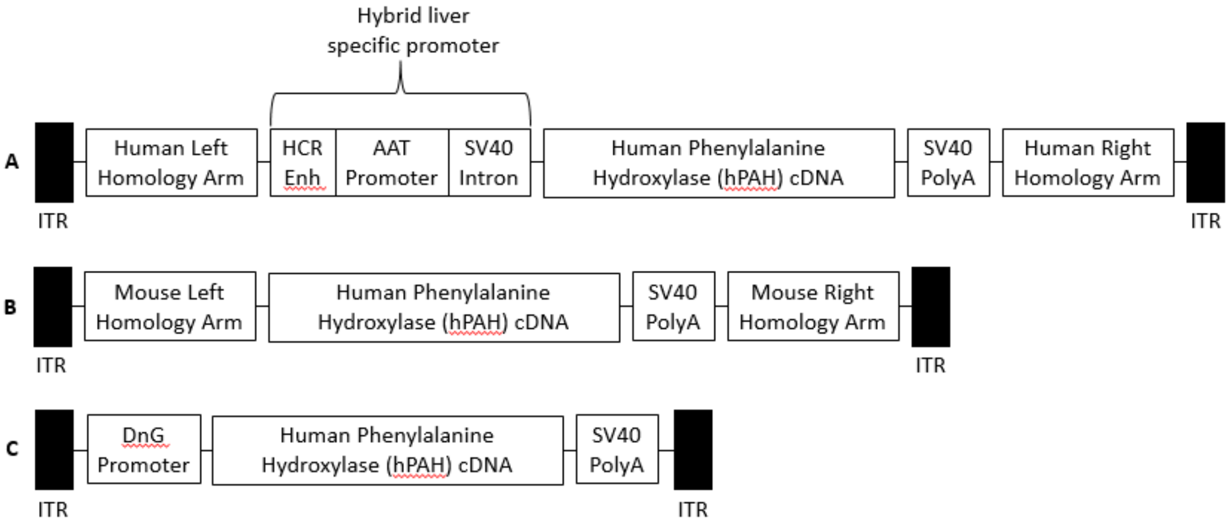


Abbreviations: AAT=alpha-1 antitrypsin; cDNA=complementary DNA; DnG=David and Goliath; Enh=Enhancer; HCR=hepatic control region; ITR=inverted terminal repeat; SV40=Simian vacuolating virus 40

A: Schematic of HMI-103 human integration construct

B: Schematic of mouse integration construct

C: Schematic of non-integrating gene transfer construct

**Supplemental Figure 2:** Schematic of splinter primer amplification

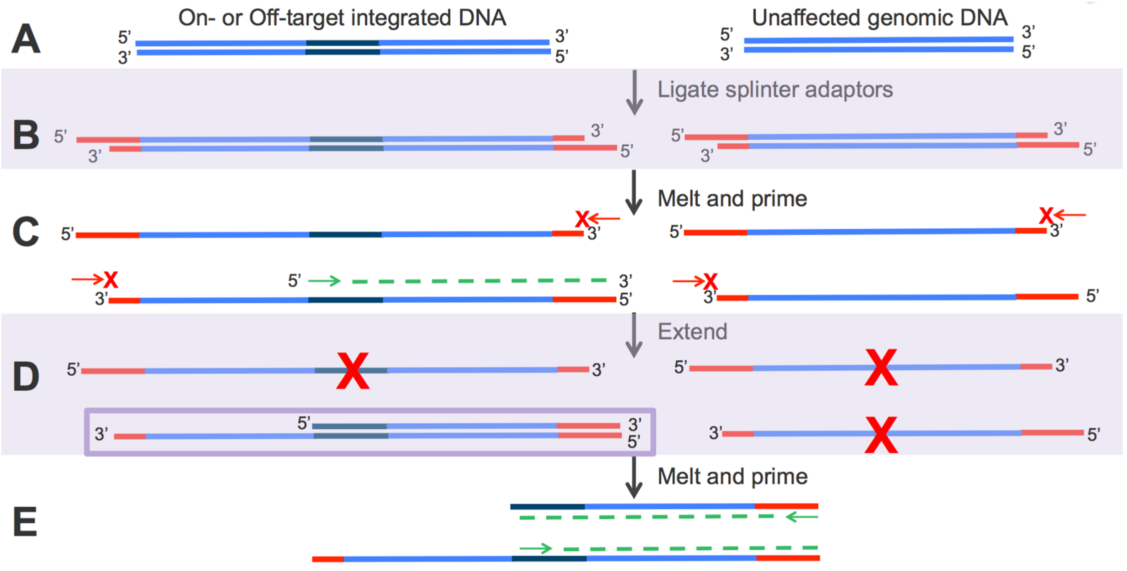


**A:** Sheared DNA

**B:** DNA with splinter adaptors ligated

**C:** First amplification – only payload primer binds

**D:** Only payload primer extends

**E:** Second amplification - both payload and one splinter primer bind and extend

**Supplemental Figure 3:** LNA Blocking of Amplification

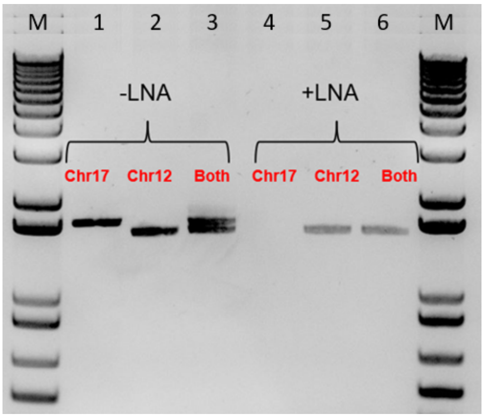


**Supplemental Figure 3:** A11 genomic DNA was amplified using two (Chr17 and Chr12) or three (Both) PCR primers. The common primer matched the CO-PAH payload, and the others were to the adjacent genomic sequence in either Chr12 or Chr17. Lanes 1-3 did not include LNAs while lanes 4-6 included LNAs that bound an AAV sequence present only in Chr17.

### Supplemental Table 1: Potential Off-Target Locations

| Region | Chromosome | Start-End | Length | Status | % ID | Site | PAH (chr12) |
|---|---|---|---|---|---|---|---|
| 1 | Chr13 | 114015389-114015438 | 50 | Tested | 81 | RASA3, intron 13 | 102917338-102917388 |
| 2 | Chr10 | 131112009-131112048 | 40 | Tested | 83 | TCERG1L, intron 9 | 102916525-102916562 |
| 3 | Chr15 | 99533687-99533726 | 40 | Removed, Repetitive | 85 | Intergenic | 102916116-102916080 |
| 4 | Chr10 | 124759578-124579616 | 38 | Tested | 85 | EEF1AKMT, 3’UTR | 102916079-102916041 |
| 5 | Chr7 | 19156075-19156119 | 45 | Tested | 62 | Intergenic | 102916560-102916691 |
| 6 | Chr10 | 71217826-71217867 | 52 | Tested | 60 | UNC5B, intron 1 | 102917367-102917405 |
